# Population Genetics of Fluctuating Selection among Individuals (FSI): a New Paradigm of Molecular Evolution

**DOI:** 10.64898/2025.12.08.693052

**Authors:** Xun Gu

**Affiliations:** Department of Genetics, Development and Cell Biology, Iowa State University, Ames, IA 50011, USA; The Laurence H. Baker Center in Bioinformatics on Biological Statistics, Iowa State University, Ames, IA 50011, USA; Program of Ecological and Evolutionary Biology, Iowa State University, Ames, IA 50011, USA

## Abstract

The FSI (fluctuating selection among individuals) theory of population genetics postulates that the fitness effect of a mutation may fluctuate among individuals with the same genotype (heterozygote or homozygote), whereas the wildtype fitness remains a constant. We conducted a comprehensive population genetic analysis and demonstrated profound impacts on the current wisdom of population genetics and molecular evolution. First, genetic drift induced by FSI, FSI-genetic drift for short, may play an important role in molecular evolution when the *N*_*e*_-genetic drift is weak, i.e., the effective population size is not very small. Second, FSI toppled the golden-standard of neutrality in molecular evolution: the substitution rate equals to the mutation rate if and only if the mutation is strictly neutral. Instead, the concept of selection-duality claims that, under FSI, slightly beneficial mutations may be subject to a negative selection, resulting in a substitution rate less than the mutations rate. Consequently, there are three types of neutrality under FSI: the fixation neutrality (substitution rate equals to mutation rate), the generic neutrality (the population mean of selection coefficient is zero), and the FSI-neutrality (the midpoint of fixation neutrality and generic neutrality). Intriguingly, our theoretical analysis shows that the null hypothesis (α = 0) of MacDonald-Kreitman (MK) test exactly corresponds to the FSI-neutrality. Rejection of the null that, say, favors the alternative α > 0, could lead to two interpretations: when the observed α is below a threshold, the selection-duality is favored; otherwise an adaptive evolution is favored. We calculated the threshold for several mammals (human, macaque, lemur and vole), and found that all those observed α values (from 3.5% to 29%) were below the threshold and so they be interpreted by the selection duality rather than adaptive evolution.

## Introduction

The ongoing central debate between neutralists and selectionists has casted some doubts about the inadequacy of population genetics that underlies the theory of molecular evolution (Kimura 1968, 1983; Hahn 2008; Hughes 2008; Wagner 2008; Nei et al. 2010; McCandlish and Stoltzfus 2014; Kern and Hahn 2018; Jensen et al. 2019; Munoz-Gomez et al. 2021; Gu 2021, 2025; Galtier 2024). de Jong et al. (2024) evaluated the validity of both neutralist and selectionist arguments, concluding that many of testable predictions were sensitive to some underlying assumptions and consistent with both viewpoints. It is therefore desirable to scrutinize existed population genetics models.

A fundamental yet implicit assumption in population genetics is that any mutation has the same fitness among those individuals with the same genotype (homozygotes or heterozygotes). That said, a neutral mutation is selectively neutral for all individuals who carry the mutation, and so forth a deleterious or beneficial mutation (Kimura 1983). However, human geneticists have well-demonstrated that mutations frequently exhibit different effects on individuals, challenging this fixed view of the mutational effects (Riordan and Nadeau 2017; Eldar et al. 2009; Raj et al. 2010; Jensen et al. 2025). Gu (2021) addressed this issue in population genetics for the first time. FSI, or fluctuating selection among individuals, refers to the phenomenon when individuals with a specific mutation exhibit a broader fitness variation, while the fitness effect of the wildtype remains a constant. For instance, a (statistically) neutral mutation could be slightly deleterious in some individuals, and slightly beneficial in others, and the neutrality should be interpreted by the means of population mean.

The biological basis of FSI is complex, which can be roughly classified into several categories. The first one is *genetic background*. Chandler et al. (2013) argued phenotypic effects of a mutation are not solely determined by the mutated gene itself, instead, they can vary depending on the genetic background of the organism. Mullis et al (2018) investigated how yeast mutations exhibited different phenotypic outcomes in individuals with different genetic backgrounds. The second category is *stochastic gene expression*, which refers to the inherent randomness (noises) in gene expression (Raj and van Oudenaarden 2007). A number of authors experimentally investigated the causes of stochastic gene expression in various organisms, such as *E*.*coli* (Elowitz et al. 2002), *Bacillus subtilis* (Ozbudak et al. 2002; Maamar et al. 2007), and *C. elegans* (Vu et al. 2015). Mutations resulting in more random fluctuations can significantly impact the fates of cells. The third category is *incomplete penetrance*, which refers to the phenomenon when individuals with a specific mutation would not always show the expected disease phenotype, both environmental exposures (Khoury 1988) or perturbations (or noises) during development (Eldar et al. 2009; Suel et al. 2007) could be the crucial factor. The final one is *the complexity of genotype-phenotype map*, which refers to the phenotypic differences observed between individuals are often due to complex genetic interactions (Dowell et al. 2010). While Lehner (2013) emphasized that the environment can significantly alter the effects of genetic mutations, Taylor and Ehrenreich (2014) suggested that higher-order interactions may play a larger role in phenotypic variation than previously thought. It should be noted that those categories are not mutually excluded (Raj et al. 2010).

In spite of the solid biological basis of FSI, the effect of FSI on population genetics and molecular evolution was largely unexplored, until recently (Gu 2021). The main reason was the presumption that the effect of FSI should be negligible because nearly-neutral mutations (slightly deleterious or slightly beneficial) may have played a major role on shaping the pattern of molecular evolution. Gu (2021) demonstrated that FSI played an important role in molecular evolution especially when the effective population size (*N*_*e*_) is not small. On the other hand, the model proposed by FSI in Gu (2021) was shown inaccurate. First, it was unclear how the FSI-model of Gu (2021) differed from existed models of fluctuating selection between generation (FSG) (Karlin and Levikson 1974; Karlin and Lieberman 1974); one may also see Gu (2024) for a recent comment. Second, the effect of FSI on the mean change of gene frequency was neglected, a controversial issue that had occurred in the study of fluctuating selection between generations (###).

This article aims at a comprehensive analysis of the Wright-Fisher diffusion model under FSI, with special references to the issue whether FSI is a challenge against the current wisdom of molecular evolution. The central intent is two-fold: (*i*) to what extent FSI can serve as a novel, important resource of genetic drift, in addition to the sampling effect of finite population size, and (*ii*) to what extent FSI can provide new insights on the neutralist-selectionist debate.

## Results

### The diffusion model of gene frequency change per generation under FSI

Consider a random mating population of a monoecioys diploid organism. In a finite population, each individual produces a large number of offspring and that exactly *N* of those survive to maturity. Let *a* and *A* be the mutant and wild-type alleles at a particular locus, respectively, whose fitness effects are additive. The FSI model postulates that the fitness effect of mutant *a* is fluctuating among individuals, whereas that of wild-type *A* remains a constant. Therefore, the relative fitness of genotype *AA, Aa*, or *aa* is, on average, given by 1, 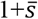 and 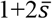, respectively; 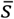 is called the mean of the selection coefficient (*s*) of mutant *a*. Let 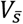 be the variance of 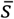 and *Var*(*s*) be the variance of *s*, respectively. Noting that the number of mutant *a* is 2*Nx*, where *x* is the frequency of mutant *a* in a generation, we obtain

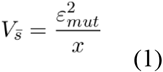

where 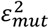 is called the FSI-coefficient; a large values means a strong FSI and *vice versa*; the subscript indicates that the FSI is induced by the mutation. In Methods, we have developed a Malthus-based model to characterize the structure of *Var*(*s*), as well as 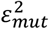.

The amount of change (*Δx*) of gene frequency per generation (where *Δt* is the generation time) under FSI can be decomposed into two independent components, that is

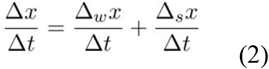

where *Δ*_*w*_*x* and *Δ*_*s*_*x* are the frequency changes caused by the Wright’s sampling and FSI, respectively. Under the Wright-Fisher model, the first and second moments of *Δ*_*w*_*x* are given by

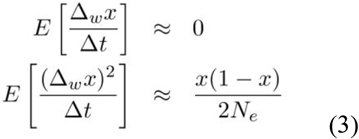

respectively, where *N*_*e*_ is the effective population size. It follows that the expected gene frequency change (per generation) with respect to FSI can be derived by

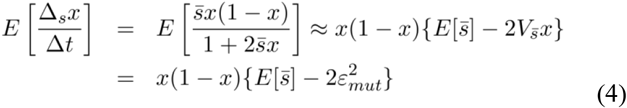

The Taylor’s expansion in Eq.(4) was carried out with respect to 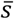, up to 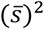, where the absolute value of 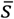 is much less than one. In the same manner, the second moment of *Δ*_*s*_*x* is given by

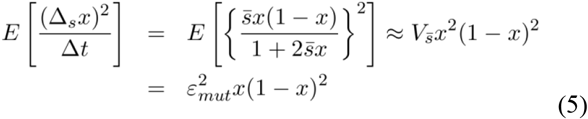

It is mathematically convenient to apply the diffusion approximation to study the Wright-Fisher model under FSI. Intuitively, the infinitesimal mean *μ*(*x*) can be approximated by 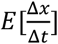, and the infinitesimal variance 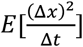. Together with Eqs.(2)-(5), we obtain

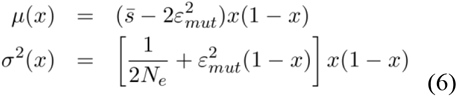

To be concise in notations, we simply use 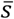 to replace 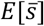 in Eq.(6). Briefly speaking, *μ*(*x*) describes the determinative factors that may influence the gene frequency change, and *σ*^*2*^(*x*) describes the random effect of genetic drifts. Noticeably, Eq.(6) indicates that FSI may have profound impacts on both selection and genetic drift. First, *μ*(*x*) is determined by the difference of selection coefficient 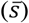 and the (two-fold) FSI coefficient 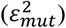. In the following analysis, the (adjusted) selection-FSI ratio (*ρ*) is more frequently used, which is defined by

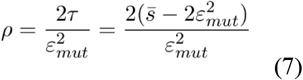

where 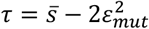. It appears that *ρ>0* indicates a positive selection, or *ρ<0* indicates a negative selection. Second, the second term of *σ*^*2*^(*x*) in Eq.(6) indicates that FSI emerges as a new resource of genetic drift. Note that *N*_*e*_ inversely measures the strength of genetic drift with respect to the Wright’s sampling process in a finite population, the *N*_*e*_-genetic drift for short. The relativity of FSI to *N*_*e*_-genetic drift can be measured by the FSI-strength: defined by 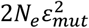, yet can be read by 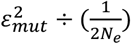. Thus, one may claim a dominant FSI-genetic drift when 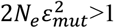, or a dominant *N*_*e*_-genetic drift when 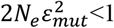. Sometimes it is more convenient to use a relative measure of FSI-strength (*F*), as given by

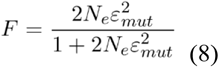

such that *F* increase from 0 at 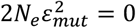 (no-FSI), which approaches to 1 when 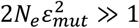.

### Substitution rate and the emergence of selection duality

The substitution rate (*λ*) plays a central role in molecular evolution (Kimura 1962). Let *v* be the mutation rate and *N* be the census population size. From the view of population genetics, the substitution rate can be defined by the amount of new mutations per generation (2*Nv*) multiplied by the fixation probability of a single mutation with the initial frequency of 1/(2*N*), based on the assumption of rare, single *de novo* mutation event. Let *u*(*p*) be the fixation probability of a mutation in a finite population, with the initial frequency *p*. Formally, the substitution rate can be written by 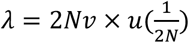. After some calculations in Methods, we obtain

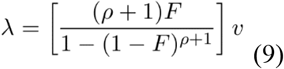

It should be noticed that *λ*>*v* (positive selection) when *ρ*>0, whereas *λ*<*v* (negative selection) when *ρ*<0. In the case of no-FSI, Eq.(9) is reduced to the well-known formula first reported by Kimura (1962).

Numerical analysis of Eq.(9) indicated that, in the case when 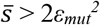 or 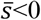, molecular evolution is mainly driven by a positive selection or a negative selection, respectively, whereas FSI only plays a marginal role. Between those two cases, an intriguing phenomenon called *selection-duality* emerges, that is, a slightly beneficial mutation falling in the range

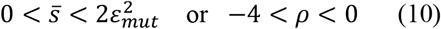

is actually subject to a negative selection, i.e., *λ*<*v* because of *ρ*<0. Apparently, this selection-duality is bounded by two types of neutralities. The low-bound one is the *generic neutrality* 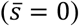, at which the mutation is neutral by the means of fitness. On the other hand, the up-bound of selection-duality is the *fixation neutrality* 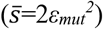, at which the substitution rate equals to the mutation rate (*λ*=*v*). The broadness of selection duality depends on the magnitude of *ε*_*mut*_^*2*^. Without FSI, i.e., *ε* _*mut*_^*2*^=0, the selection duality vanishes as the generic neutrality and the fixation neutrality merge onto the conventional strict neutrality. In addition, we notice that the middle-point of selection-duality at 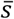, called *FSI-neutrality*, may play a pivotal role in the new population genetics theory of molecular evolution, as will be shown later.

For illustration, we derived the substitution rate in several special cases. In addition to the fixation neutrality (*ρ*=0), FSI-neutrality (*ρ*=-2) and generic neutrality (*ρ*=-4), we also calculated the cases of *ρ*=-1 and *ρ*=-3 to represent the case of selection-duality. These results are summarized as follows

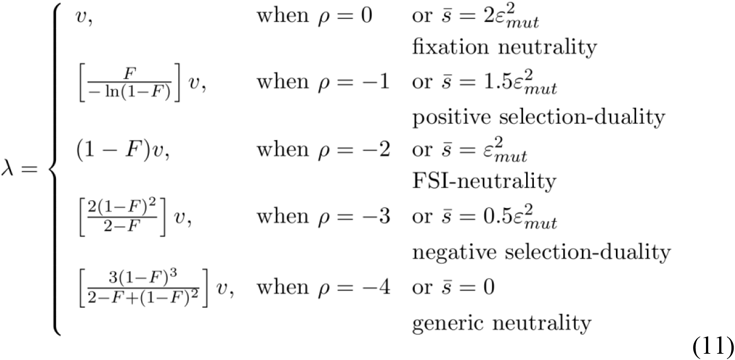

As shown by Fig.1, the substitution rate (λ) under selection-duality decreases considerably as the FSI-strength (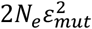 or *F*) increases. In the worst situation, the substitution rate at generic neutrality 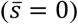 is only 21% or 8.1% of the mutation rate when 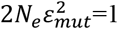 or 2, respectively. Invoking Ohta’s criterion, one may conclude that genetic neutral mutations behalf as nearly-neutral mutations when 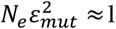. In other words, selection-duality mutations may be dominant during the sequence evolution.

**Fig. 1.**
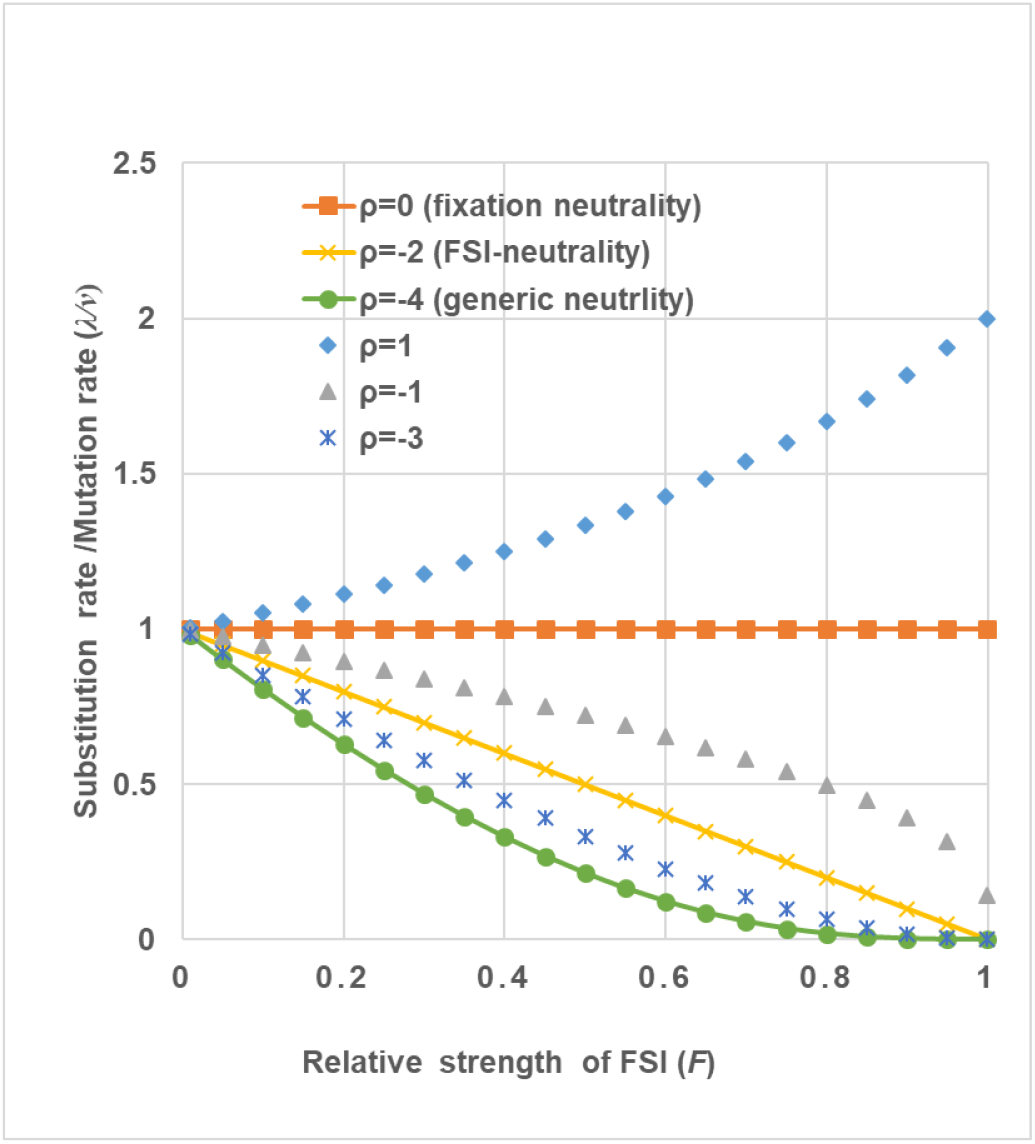
The impact of FSI on the substitution rate under selection duality. Three types of neutrality are presented: the fixation neutrality, FSI-neutrality and generic neutrality. In addition, the case of *ρ*=-1 for illustrating the theme of positive selection-duality 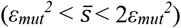, and the case of *ρ*=-3 for the theme of negative selection-duality 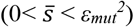, as well as the case of *ρ*=1 for beneficial mutations. Selection-duality is defined by the between the solid line of fixation neutrality (*ρ*=0) and that of generic neutrality (*ρ*=-4).

### Mean fixation time (generations) of a new mutation under selection-duality

How many generations it will take for a mutant to be fixed in a finite population is a fundamental problem of population genetics. To this end, one may have to trace a particular mutant and study the average number of generations at which the frequency of the allele become 1 (fixed). Let *T*_*fix*_(*p*) be the mean fixation time of a mutation, given the initial frequency *p*. There are many approaches to deriving *T*_*fix*_(*p*) fixed, yet all of them gave virtually the same result; one may see Crow and Kimura (1970) or Ewens (2004) for discussion. Here we adopt the method of Kimura and Ohta (1969) for simplicity (see Methods). In particular, we are interested in the mean fixation time of a new mutation arising in the population, with the initial frequency *p*=1/(2*N*), where *N* is the census population size. Except for a very small population size, the mean fixation time for a nearly arising mutation can be well approximated by letting *p* → 0; one may write

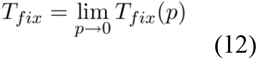

Following the procedure formulated in Methods, we computed *T*_*fix*_ in serveral special cases of selection-duality that are analytical; they are, respectively, given by

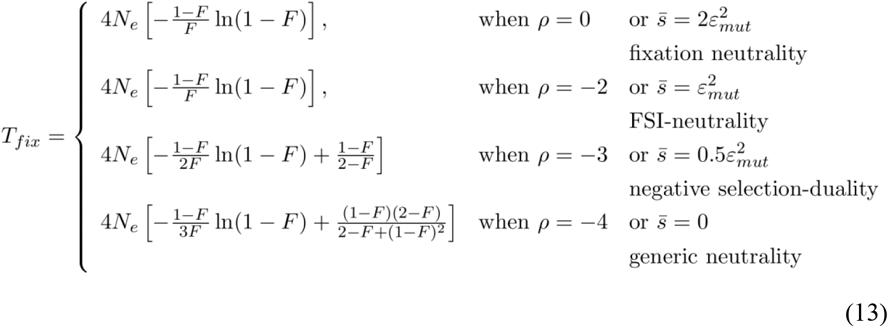

It should be noticed that, in the case of no-FSI, i.e., the relative FSI-strength *F*=0, the mean fixation time approaches to a well-known result *T*_*fix*_ = 4*N*_*e*_, i.e., a neutral mutation would take, on average, 4*N*_*e*_ generations to be fixed in a population. As FSI becomes strong, *T*_*fix*_ can be shortened considerably. Unexpectedly, Eq.(13) reveals that *T*_*fix*_ is precisely identical between the case of fixation neutrality (*ρ*=0) and FSI-neutrality (*ρ*=-2). Fig.2 represents the plotting of *T*_*fix*_/4*N*_*e*_ against the relative FSI-strength (*F*) in the range of selection duality. It appears that the mean fixation time required for a selection-duality mutation only marginally depends on the ratio *ρ*=0, which, overall, could be much shorter than 4*N*_*e*_ generations when FSI is strong.

**Fig. 2.**
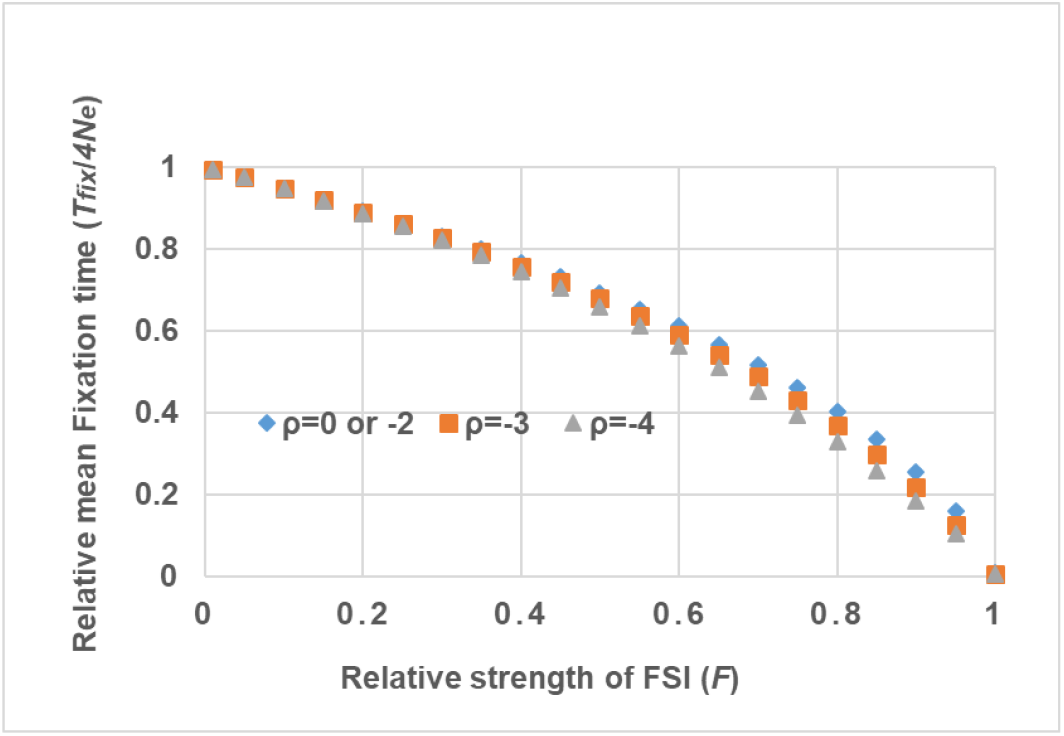
Mean fixation time (*T*_*fix*_) under fixation neutrality, FSI-neutrality and generic neutrality, respectively. The ratio of mean fixation time (*T*_*fix*_) to 4*N*_*e*_ (the mean fixation time in the strict neutrality), plotting against the relative strength of FSI (*F*), revealing a reduced (*T*_*fix*_) with the increasing of FSI strength.

On the other hand, the average number of generations for a new mutation to be lost from the population (*T*_*loss*_) can be obtained similarly. For instance, an identical formula of *T*_*loss*_was derived in the case of *ρ*=0 or *ρ*=-2, given the small initial frequency *p*=1/(2*N*), that is,

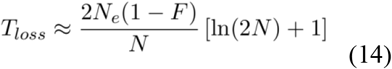

Two properties are mentioned. First, when *F*=0 (no FSI), *T*_*loss*_ converges to the classical result in the theme of selection-duality. And second, FSI accelerates the loss process of a selection-duality mutation.

### Steady-state spectrum of allele frequency

We assume that, at the nucleotide site level, the mutation rate is so low that a population (with a finite population size) is almost always monomorphic or polymorphic for two types, the mutant type and the wild type. Reversible mutation is virtually negligible while they are polymorphic, suggesting that the two-allele theory of population genetics applies. Under this scenario, each nucleotide may mutate independently and the mutation type may increase or decrease in frequency. At equilibrium when the effects of mutation, selection and genetic drift are balanced, one may expect that the frequency of mutant sites reaches a steady-state spectrum, denoted by Φ(*x*) (Sawyer and Hartl 1992). A general formula of Φ(*x*) under FSI has been derived (see Methods); some typical cases under selection duality of FSI are presented by

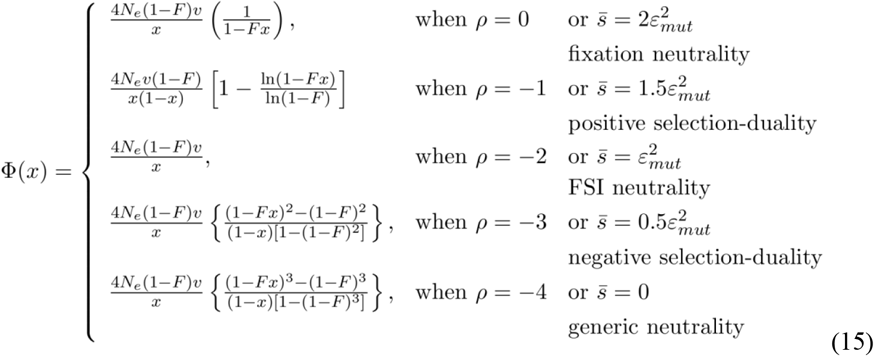

As expected, in the case of no FSI, i.e., *F*=0, Φ(*x*) under selection-duality would converge to the classical result of strict neutrality, that is,

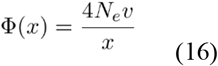

Fig.3 presents the plotting of Φ(*x*)/4*N*_*e*_*v* against the allele frequency (*x*), under various values of relative FSI-strength (*F*); panel A for fixation neutrality, panel B for FSI-neutrality, and panel C for generic neutrality. In particular we noticed that Φ(*x*) under FSI-neutrality (*ρ*=-2) in panel B has virtually the same shape as Φ(*x*) under the strict neutrality, with a rescaled factor 4*N*_*e*_*v*(1 − *F*). Meanwhile, Φ(*x*) under the fixation neutrality in panel A reveals an excess of intermediate or high allele frequency, usually considered as a sign of positive selection. By contrast, Φ(*x*) under the generic neutrality in panel C reveals an excess of low allele frequency, a sign usually considered as negative selection. It is therefore rational to divide selection-duality into two regions: the *positive selection-duality* between fixation neutrality and FSI-neutrality, i.e., −2<*ρ*<0, and the *negative selection-duality* between FSI-neutrality and generic neutrality, i.e., −4<*ρ*<-2.

**Fig. 3.**
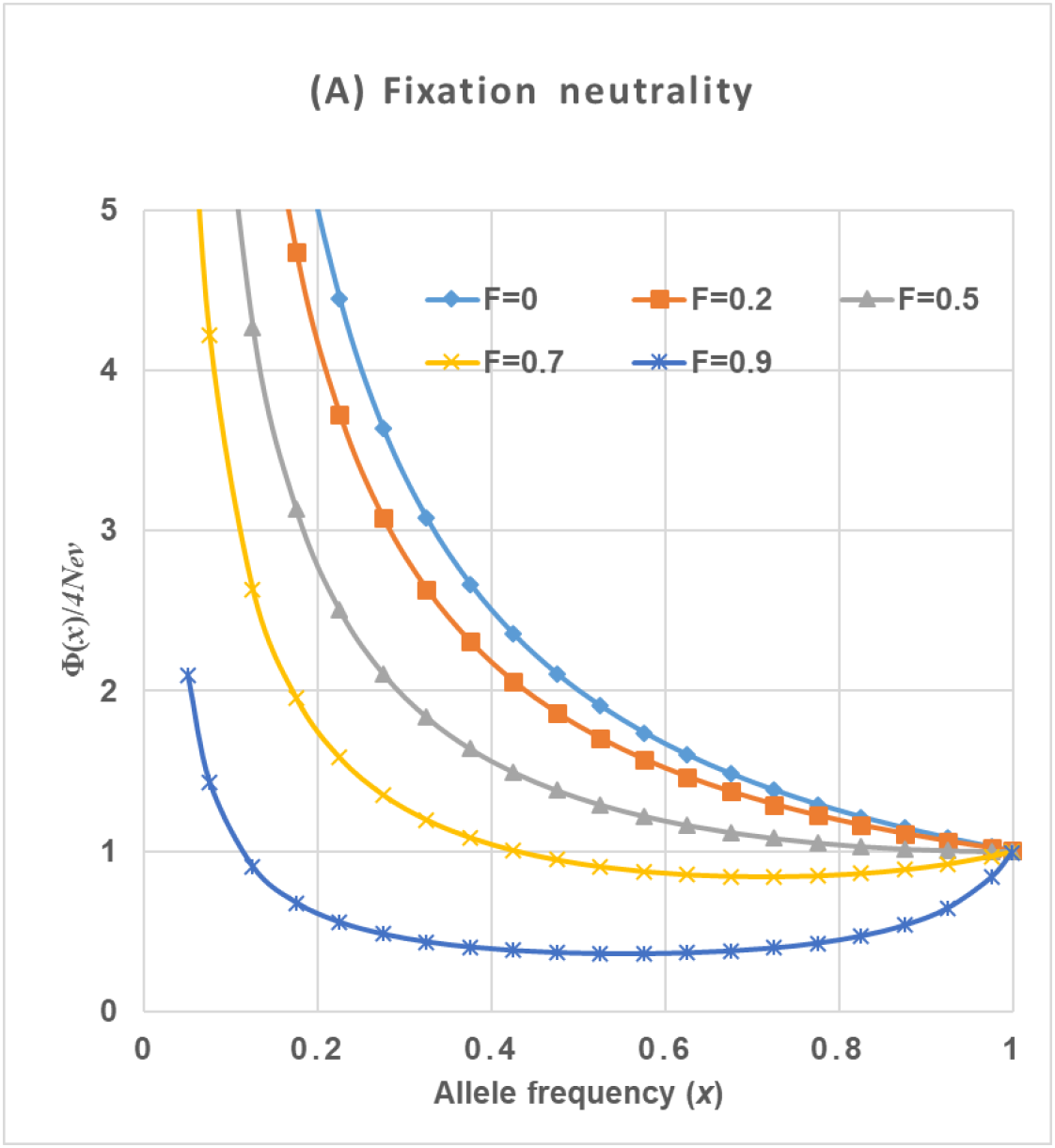
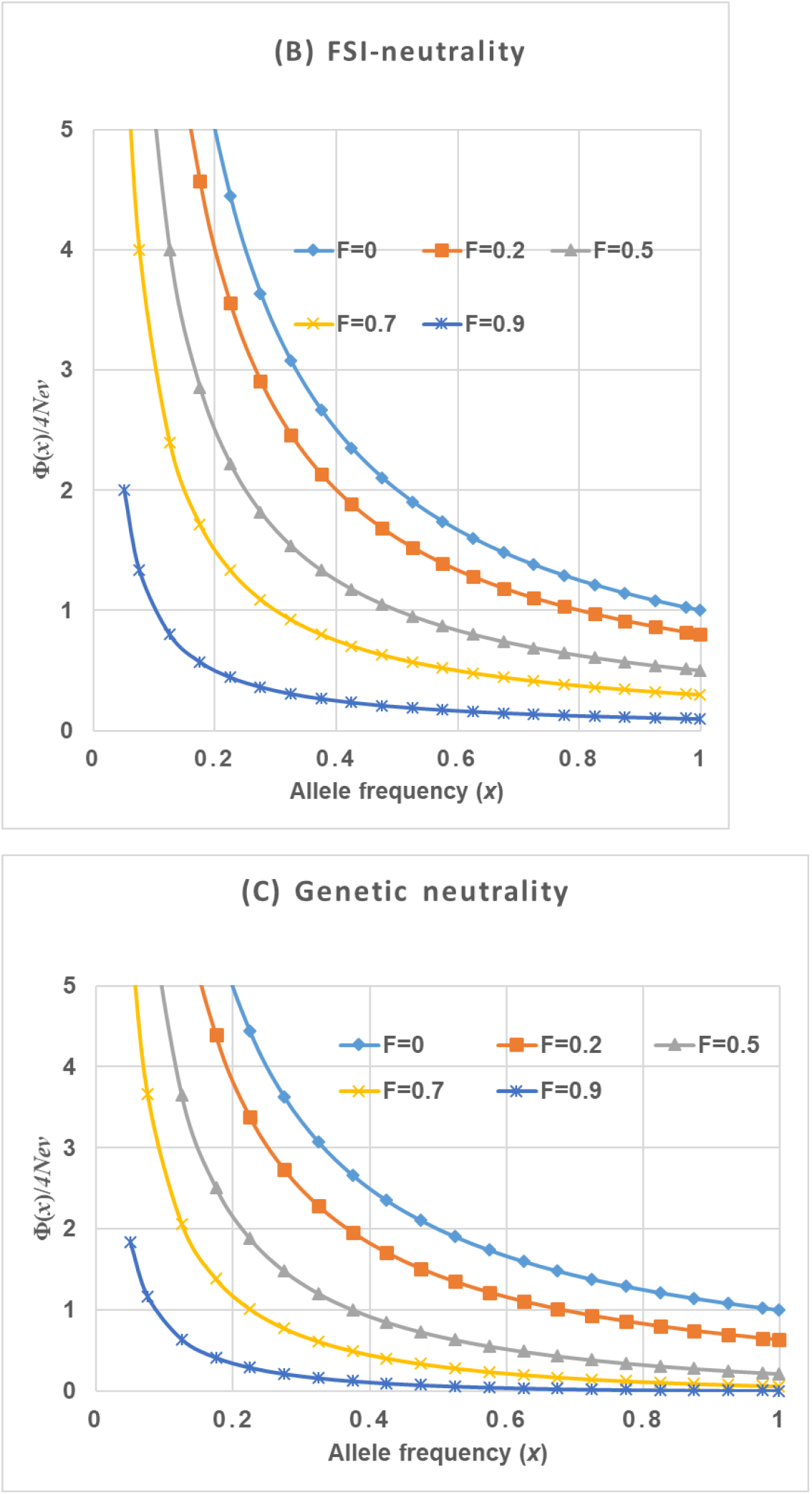
Plotting of Φ(*x*)/4*N*_*e*_*v* against the allele frequency (*x*), under various values of relative FSI-strength (*F*). (A) Φ(*x*) under the fixation neutrality reveals an excess of intermediate or high allele frequency, usually considered as a sign of positive selection. (B) Φ(*x*) under FSI-neutrality (*ρ*=-2) has virtually the same shape as Φ(*x*) under the strict neutrality, with a rescaled factor 4*N*_*e*_*v*(1 − *F*). (C) Φ(*x*) under the generic neutrality reveals an excess of low allele frequency, a sign usually considered as negative selection. Thus, selection-duality is divided into two regions: the *positive selection-duality* between fixation neutrality and FSI-neutrality (−2<*ρ*<0), and the *negative selection-duality* between FSI-neutrality and generic neutrality (−4<*ρ*<-2).

When Φ(*x*) is derived, it is straightforward to calculate the genetic diversity, defined by the expected heterozygosity at a nucleotide site, that is,

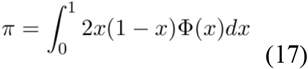

While the general result can be seen in Methods, the genetic diversity in the case of fixation neutrality, FSI-neutrality or generic neutrality is given by

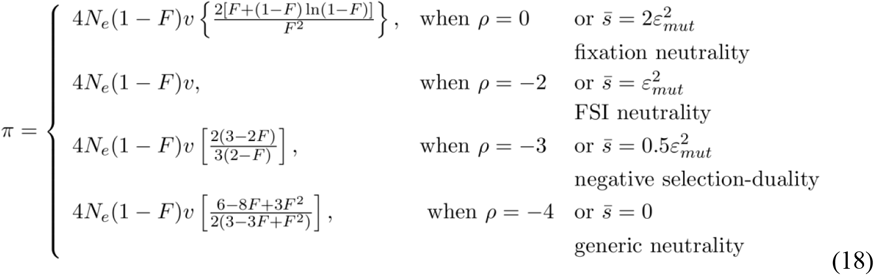

respectively.

### FSI-neutrality served as null hypothesis of molecular evolution

The excess ratio of sequence divergence (α) between species compared to the genetic diversity within population has played a pivotal role in the neutrality-selection debate (Hahn 2008; Kern and Hahn 2018; Jensen et al. 2019; Galtier 2024; de Jong et al. 2024). An intriguing question is how FSI could affect the excess ratio α. Let *R* be the scaled diversity-divergence ratio, defined by the ratio of genetic diversity (π) (scaled by 4*N*_*e*_*v*) to the substitution rate (λ) (scaled by *v*). According to Eq.(9) and Eq.(17), one can show

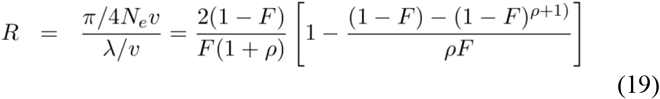

It follows that the theoretical value of α is given by α = 1 − *R*. We further present the excess ratio α with respect to the relative FSI-strength (*F*) under various selection scenarios, that is,

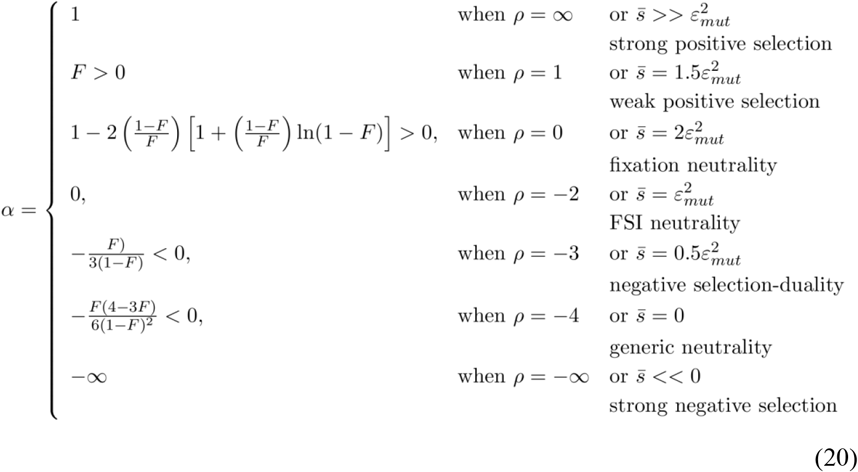

Eq.(20) provides a theoretical foundation to interpret the null hypothesis of MK test. Under FSI, the null hypothesis *α* = 0 means FSI-neutrality (*ρ* = −2), i.e., the mean selection coefficient satisfies 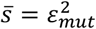, which is slightly beneficial. Rejection of this null hypothesis suggested the alternative, say, α > 0. As illustrated by Fig.4, under FSI it could be explained by either positive selection-duality (−2 < *ρ* < 0), or positive selection (ρ > 0) that indicates adaptation. Intuitively, one may calculate the theoretical *α*-index at the fixation neutrality (*ρ* = 0) to distinguish between these two possibilities. Similarly, in the case of *α* < 0, one may calculate the α-index value at the generic neutrality (*ρ* = −4) to distinguish between negative selection-duality (−4 < *ρ* < −2) and the purifying selection (ρ < −4 or 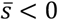).

**Fig. 4.**
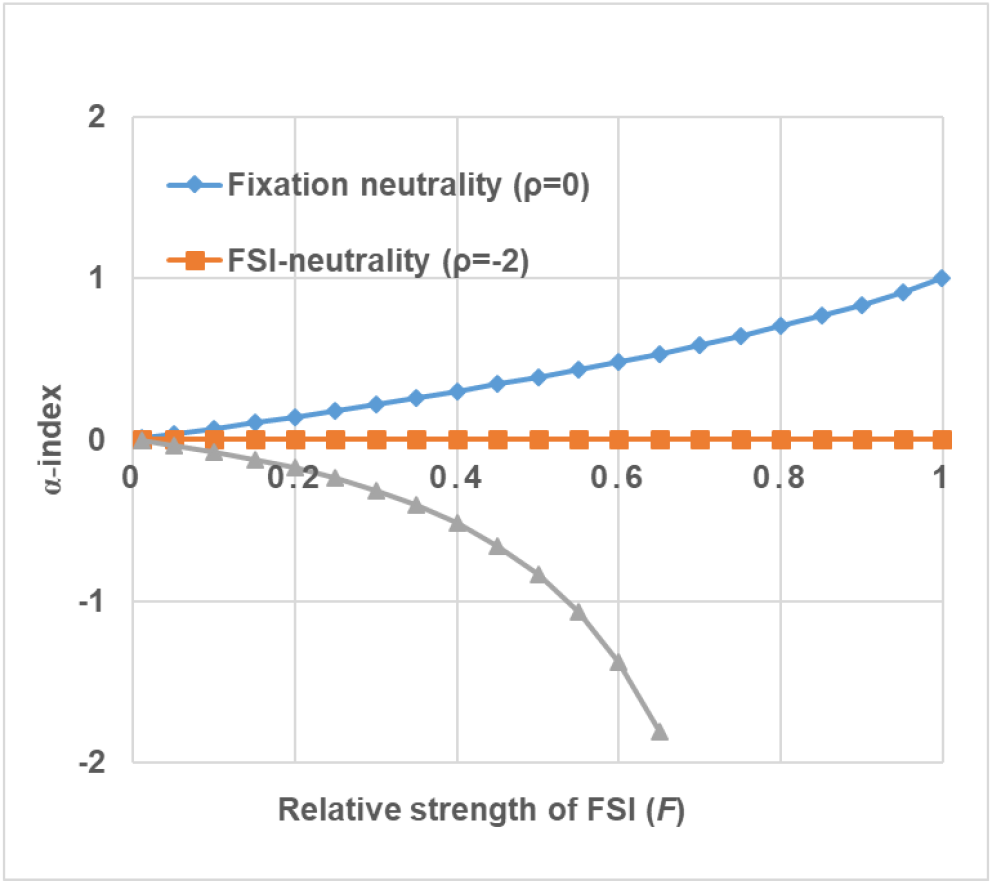
Plotting of the theoretical α-index against various values of relative FSI-strength (*F*). Under FSI, the case of α > 0 could be explained by either positive selection-duality (−2 < ρ < 0), or the conventional positive selection (ρ > 0) that may indicate an adaptive evolution. The theoretical α-index at the fixation neutrality (ρ = 0) is to distinguish between them. In the case of α < 0, the theoretical α-index value at the generic neutrality (ρ = −4) is to distinguish between negative selection-duality (−4 < ρ < −2) and the conventional purifying selection (ρ < −4 or 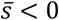).

Galtier et al. (2016) reported a genome-wide analysis in metazoans, a genome-wide set of protein-coding genes in 44 pairs including 18 vertebrate and 26 invertebrate pairs. Each pair consisted of two closely-related species to estimate the synonymous distances (*d*_*S*_) and the nonsynonymous distance (*d*_*N*_), respectively. Moreover, the synonymous genetic diversity (π_*S*_) in the focal species of each specie pair was used as a proxy of the effective population size. After extending the nearly-neutral of Ohta (1973; Ohta 1993), in a separate research Gu (2025) predicted the relative FSI-strength (*F*) for each species under study. Table 1 presents the analysis of four mammalian species for illustration; in each case we calculated the theoretical α-indexes for the fixation neutrality and the generic neutrality, respectively, and found the observed α-indexes are all within the region of (positive) selection-duality (Table 1). The increase of observed α-indexes with the synonymous diversity can be simply explain by the reduced *N*_*e*_-genetic drift as the result of increased effective population size.

**Table 1.**
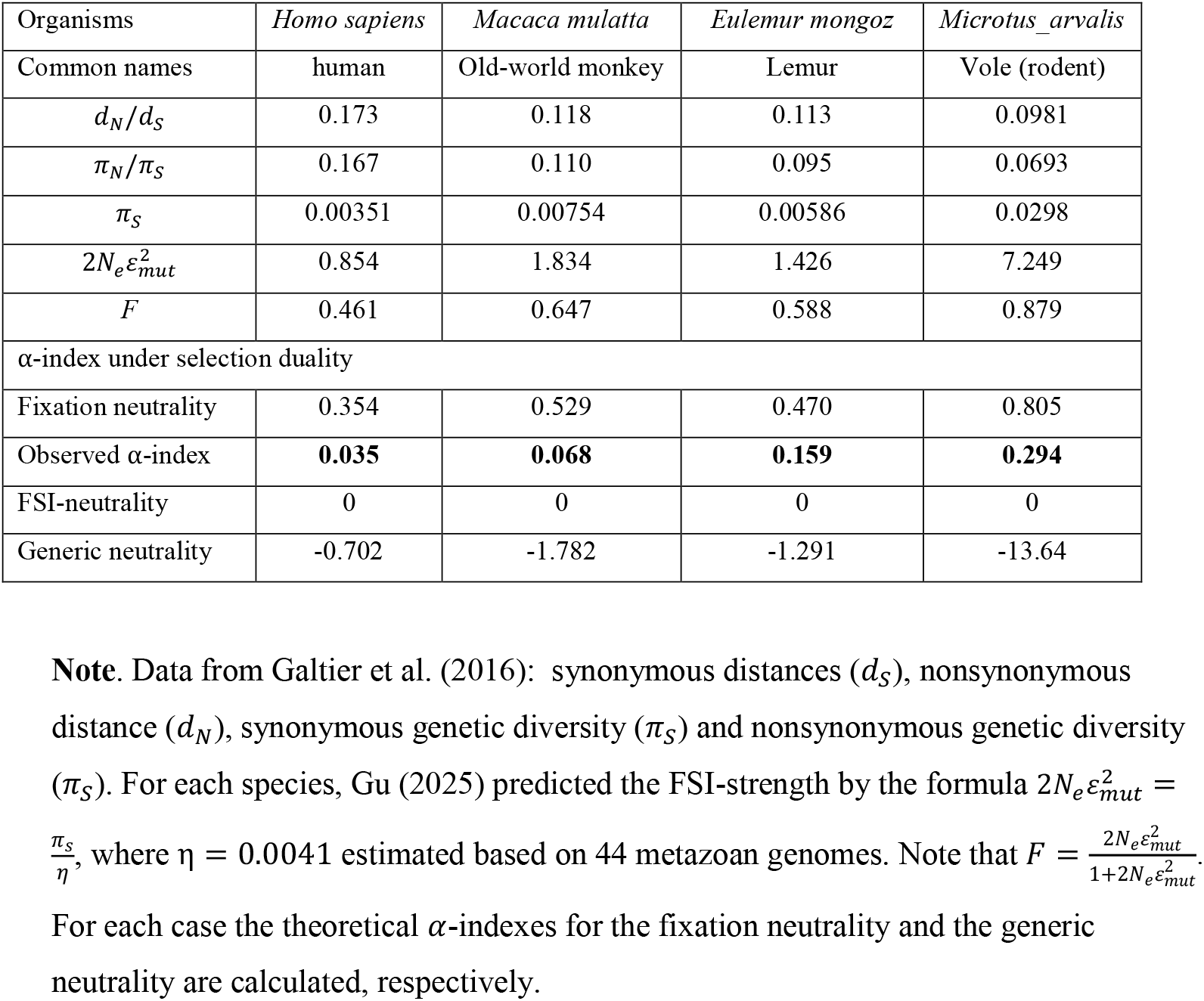
Some examples for the α-index under selection duality.

## Discussion

The FSI (fluctuating selection among individuals) theory postulates that the fitness effect of a mutation may fluctuate among individuals with the same genotype (heterozygote or homozygote), whereas the fitness of the wildtype remains a constant. The analysis reported in this paper showed several profound impacts on the current wisdom of population genetics. First, genetic drift induced by FSI may play an important role in molecular evolution when the effective population size is not very small. Second, FSI toppled the golden-standard of neutrality in molecular evolution: the substitution rate equals to the mutation rate if and only if the mutation is strictly neutral. The concept of selection-duality claims that, under FSI, slightly beneficial mutations may be subject to a negative selection, resulting in a substitution rate that is less than the mutations rate. One may anticipate that FSI is intrinsically related to the evolution and maintenance of phenotypic plasticity that may imply local adaptation (Sommer 2020; Lee et al. 2022; Wang et al. 2023). Moreover, the effect of shaping eukaryotic genomes by genetic drift (Lynch et al. 2011) should be extended to FSI.

Overall, FSI-theory shed some lights on resolving the long-term neutralist-selectionist debate. As shown by a separate work (Gu 2025), selection duality mutations may dominate the pattern of molecular evolution between species, compatible with the observed inverse relationship between the substitution rate and the (log-of) effective population size. Under the null hypothesis of FSI-neutrality, most statistical tests about neutrality in population genetics are legitimate, but the interpretation needs to be revised appropriately. We use the MK test for illustration. First, the null hypothesis (α = 0) does not mean a strict neutrality; rather, it represents a specific status of balance between (statistically beneficial) selection and FSI. Second, rejection of the null with, say, the alternative α > 0, there are two possibilities: when α is below a threshold, the selection-duality would be favored; otherwise when α exceeds the threshold, an adaptive evolution is favored. We calculated the threshold for several mammals and found that the observed α values ranged from 3.5% to 29%, all of them were below the threshold and so can be interpreted by the selection duality (Table 1).

An unexpected result is that the mean fixation time is, roughly, in the same magnitude for selection-duality mutations, which is much shorter than that of a strictly neutral mutation. This fact provides a new explanation for the ‘fixation’ sweeps at linked sites. Tentatively, we consider a simple hard fixation sweep that had happened recently, that is, some strictly neutral sites that are tightly linked (no crossing-over) to the site that is undergoing a fixation process of a mutation. One interesting question is, if this fixing mutation is strictly neutral, to what extent the genetic diversity at linked sites can be reduced after the fixation process. Since the mean fixation time of a strictly neutral mutation is 4*N*_*e*_, the expected diversity at a tightly liked sites is accumulated up to 4*N*_*e*_*v* (*v* is the mutation rate). That is, fixation of a strictly neutral mutation would not result in any genetic diversity reduction at linked sites.

Hence, selective sweep has been widely invoked to account for a dramatic reduction of genetic diversity in some genome regions, which can be intuitively explained by following the logic of Gillespie (2000; 2001). For instance, when *N*_*e*_*s* = 2.5, the mean fixation time is about half that of a strict neutral mutation, predicting a 50% reduction of genetic diversity at linked sites. The challenge against the selective sweeps is the lack of empirical evidence for the pervasive genome-wide adaptations. Instead, the impact of selection-duality sweep only depends on the FSI-strength: For 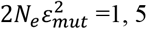, or 20, the gene diversity at liked strictly neutral sites would be reduced to 69%, 36% or 15%, respectively.

We emphasize that election-duality would be not rule out other explanations in the neutralist-selectionist debate, such as demographic history versus adaptive evolution. Yet, the notion of selection-duality does explain the pattern of negative selection between species, the excess of sequence divergence versus diversity (α > 0 in MK test) and the genetic reduction by fixation under a universal framework.

## Methods

### The Malthus model of FSI

Under this additive genetic model and the Hardy-Weinberg equilibrium, it is known that the diploid Wright-Fisher model can be simplified to a simple two-allele system. Let *a* and *A* be the mutant and wild-type alleles at a particular locus, respectively, and *x* be the frequency of mutant *a* in one generation. Let *Z*_*A*_ or *Z*_*a*_ be the number of chromosomes produced by one parent *A*-chromosome or *a*-chromosome, respectively. FSI postulates that *Z*_*a*_ is random variable, whereas *Z*_*A*_ is a constant. Under a finite population, we further denote *z*_*a*_ to be the population mean of *Z*_*a*_, and 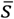 to be the population mean of selection coefficient of mutation-*a*, respectively. Under this Malthus model, we have

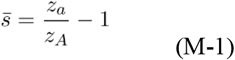

We try to derive Eq.(1), the variance of 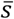, denoted by 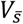. When the random fluctuation in selective coefficient is mainly due to the fluctuation in genetic or epigenetic backgrounds that might be individual-specific, the effect of FSI on the change of allele frequency in each generation should be determined by the variance of its population mean (*z*_*a*_). Let *V*(*Z*_*a*_) be the variance of *Z*_*a*_. Since the number of *a*-chromosomes is expected to be 2*N*_*e*_*x*, where *x* is the frequency of mutation and *N*_*e*_ is the effective population size, we have

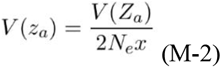

By Eq.(M-1) it is straightforward that the variance of 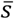 is given by

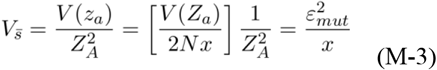

which is inversely related to the frequency of mutation-*a*, as first speculated by Ohta (1973).

### Derivation of the substitution rate

We first calculate the function *G*(*x*) as follows

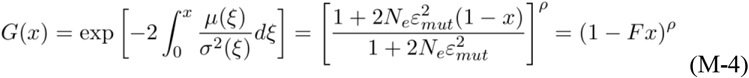

Let *u*(*p*) be the fixation probability of a mutation in a finite population, with the initial frequency *p*. It follows that the fixation probability can be calculated by

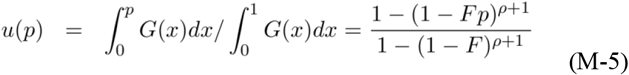

The formula of substitution rate can be concisely written as follows

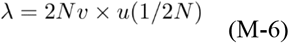

We try to solve Eq.(M-6) under the approximation of very small initial frequency (*p* → 0) in Eq.(M-5), where the fixation probability is approximately given by

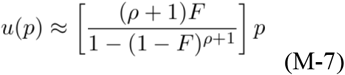

Together with Eq.(M5) and (M6), we obtained the general formula of the substitute rate of Eq.(9). In the case of no FSI, i.e., *ε* _*mut*_^*2*^=0, one can verify that 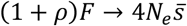, and 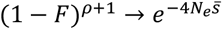, respectively, resulting in the well-known formula of Kimura (1962), i.e.,

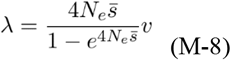

### Derivation of the mean fixation time

Let *T*_*fix*_(*p*) be the mean fixation time of a mutation, given the initial frequency *p*. Kimura and Ohta (1969) showed that *T*_*fix*_(*p*) can be calculated by

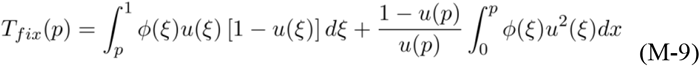

where *u*(*x*) is the fixation probability, and ϕ(*x*) by

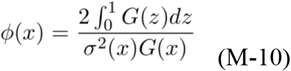

We are particularly interested in the mean fixation time of a new mutation with a very small initial frequency *p*=1/(2*N*), where *N* is the census population size. Hence, the mean fixation time of a new mutation can be approximated by *p* → 0, that is

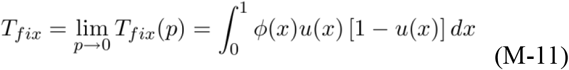

### Derivation of the mean loss time

In the same manner, the mean loss time of a mutation can be computed by

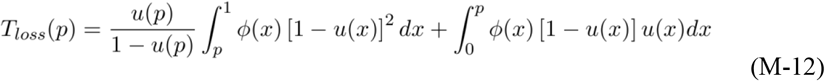

For a new mutation with a very low initial frequency, one may claim

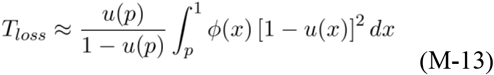

### The steady-state spectrum of allele frequency

We assume that, at the nucleotide site level, the mutation rate is so low that a population (with a finite population size) is almost always monomorphic or polymorphic for two types, the mutant type and the wild type. Reversible mutation is virtually negligible while they are polymorphic, suggesting that the two-allele theory of population genetics applies. In this case each nucleotide may mutate independently and the mutation type may increase or decrease in frequency. At equilibrium (the steady-state condition) when the effects of mutation, selection and genetic drift are balanced, one may expect that the frequency of mutant sites reaches a steady-state spectrum. It has been shown that the general formula of the steady-state is given by

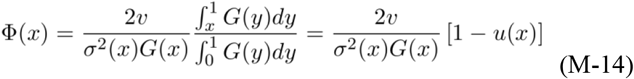

where *v* is the mutation rate.

## References

Chandler et al. (2013) Does your gene need a background check? How genetic background impacts the analysis of mutations, genes, and evolution. Trends Genet

Crow J, Kimura M. 1970. An Introduction to Population Genetics Theory. Minneapolis (MN): Burgess Publishing Company.

de Jong et al. (2024) Moderating the neutralist–selectionist debate: exactly which propositions are we debating, and which arguments are valid? Biological Reviews

Dowell et al. (2010) Genotype to phenotype: a complex problem. Science 328:469.

Elowitz et al. (2002) Stochastic Gene Expression in a Single Cell. Science

Eldar A et al. (2009) Partial penetrance facilitates developmental evolution in bacteria. Nature 460:510–514.

Ewens WJ. 2004. Mathematical Population Genetics: theoretical Introduction. Vol. 1. New York (NY): Springer.

Galtier, N. (2016). Adaptive protein evolution in animals and the effective population size hypothesis. PLoS Genetics.

Galtier (2024) Half a Century of Controversy: The Neutralist/Selectionist Debate in Molecular Evolution. GBE

Gillespie JH (2000) The neutral theory in an infinite population. Gene 261:11–18. 32.

Gillespie JH (2001) Is the population size of a species relevant to its evolution? Evolution 55:2161–2169.

Gu, X (2021) Random penetrance of mutations among individuals: a new type of genetic drift in molecular evolution. Phenomics

Gu, X (2024) Models of fluctuating selection between generations: a solution for the theoretical inconsistency. Journal of Molecular Evolution.

Gu, X (2025) Fluctuating Selection among Individuals (FSI) as a Novel Genetic Drift in Molecular Evolution. BioRxiv.

Hahn MW (2008) Toward a selection theory of molecular evolution. Evolution 62:255– 265.

Hughes, A. L. (2008). Near-neutrality: the leading edge of the neutral theory of molecular evolution. Annals of the New York Academy of Sciences

Jensen et al. (2019) The importance of the Neutral Theory in 1968 and 50 years on: A response to Kern and Hahn 2018. Evolution 73:111–114.

Jensen et al. (2025) Genetic modifiers and ascertainment drive variable expressivity of complex disorders. Cell

Karlin and Levikson. 1974. Temporal fluctuations in selection intensities: Case of small population size. Theoret. Popul. Biol. 6:383–342

Karlin, S, and U. Lieberman, 1974 Random temporal variation in selection intensities: case of large population size. Theoret. Popul. Biol. 6: 355–382

Kern AD, Hahn MW (2018) The Neutral Theory in Light of Natural Selection. Mol Biol Evol 35:1366–1371.

Kimura M (1962) On the probability of fixation of mutant genes in a population. Genetics 47:713–719

Kimura M (1968) Evolutionary rate at the molecular level. Nature 217:624–626.

Kimura M (1983) The Neutral Theory and Molecular Evolution. Cambridge University Press, New York

Kimura M, Ohta T. 1969. The average number of generations until fixation of a mutant gene in a finite population. Genetics. 61(3): 763. doi:10.1093/genetics/61.3.763

Khoury MJ, Flanders WD, Beaty TH (1988) Penetrance in the presence of genetic susceptibility to environmental factors.

Lee et al. (2022) Evolution and maintenance of phenotypic plasticity. Biosystems.

Lehner B (2013) Genotype to phenotype: lessons from model organisms for human genetics. Nat Rev Genet 14:168–178.

Lynch M et al. (2011) The repatterning of eukaryotic genomes by random genetic drift. Annu Rev Genomics Hum Genet 12:347–366.

Maamar H, Raj A, Dubnau D (2007) Noise in gene expression determines cell fate in Bacillus subtilis. Science 317

McCandlish and Stoltzfus (2014) Modeling Evolution Using the Probability of Fixation: History and Implications. The Quarterly Review of Biology

Mullis et al (2018) The complex underpinnings of genetic background effects. Nat Commun 9:3548.

Munoz-Gomez et al (2021). Constructive neutral evolution 20 years later.

JME Nadeau JH (2001) Modifier genes in mice and humans. Nat Rev Genet 2:165–174.

Nei, M., Suzuki, Y. & Nozawa, M. (2010). The neutral theory of molecular evolution in the genomic era. Annual Review of Genomics and Human Genetics

Ohta T (1973) Slightly deleterious mutant substitutions in evolution. Nature 246:96–98.

Ohta T (1993) An examination of the generation-time effect on molecular evolution. Proc Natl Acad Sci U S A 90:10676–10680.

Ozbudak et al. (2002) Regulation of noise in the expression of a single gene. Nat Genet

Raj A et al. (2010) Variability in gene expression underlies incomplete penetrance. Nature 463:913–918.

Raj A, van Oudenaarden A (2008) Nature, nurture, or chance: stochastic gene expression and its consequences. Cell

Riordan JD, Nadeau JH (2017) From Peas to Disease: Modifier Genes, Network Resilience, and the Genetics of Health. Am J Hum Genet 101:177–191.

Sawyer, S. A., and D. L. Hartl, 1992 Population genetics of polymorphism and divergence. Genetics 132:1161–1176.

Sommer (2020) Phenotypic Plasticity: From Theory and Genetics to Current and Future Challenges. Genetics

Süel (2007) Tunability and noise dependence in differentiation dynamics. Science

Taylor MB, Ehrenreich IM (2014) Genetic interactions involving five or more genes contribute to a complex trait in yeast. PLoS Genet

Vu V et al (2015) Natural Variation in Gene Expression Modulates the Severity of Mutant Phenotypes. Cell

Wagner, A. (2008). Neutralism and selectionism: a network-based reconciliation. Nature Reviews Genetics

Wang et al. (2023) Reproductive variance can drive behavioral dynamics. PNAS

